# Coevolution and synchronized evolutionary rates in aphid dual endosymbiosis

**DOI:** 10.64898/2026.05.09.722923

**Authors:** Jess Rouïl, Alejandro Manzano-Marín, Anne-laure Clamens, Johannes Tavoillot, Corinne Cruaud, Valerie Barbe, Armelle Coeur d’acier, Emmanuelle Jousselin

## Abstract

Many insects rely on obligate bacterial endosymbionts for essential nutrients. However, long-term endosymbiosis drives genome erosion, frequently resulting in the acquisition of additional symbionts that complement or replace ancestral partners. In aphids, the primary nutritional symbiont *Buchnera aphidicola* can be supplemented by a co-obligate symbiont, most commonly *Serratia symbiotica*. The long-term evolutionary trajectory of this newly acquired symbiont and its impact on *Buchnera* remain unknown. Here, we assembled host mitochondrial and endosymbiont genomes from thirteen aphid species belonging to a clade in which *Serratia* has functioned as a co-obligate symbiont for approximately 25 million years. Phylogenomic analyses reveal that *Serratia* has undergone extensive genome reduction followed by strict codivergence with aphids and *Buchnera*. Bayesian molecular dating shows that substitution rates in *Serratia* and *Buchnera* are tightly correlated and fall within the range previously reported for *Buchnera*. Genome-wide analyses indicate pervasive purifying selection in both symbionts. Homologous host-provisioning genes retained in both symbionts did not exhibit elevated evolutionary rates. However, unlike other functional categories, their dN/dS values were not correlated between symbionts, suggesting that they no longer evolve under shared selective regimes, consistent with progressive metabolic specialization. Intraspecific patterns of polymorphism and phylogenies mirror macroevolutionary patterns across the clade. Fluorescence in situ hybridization shows that *Serratia* and *Buchnera* occupy distinct bacteriocytes, suggesting that similar demographic processes may contribute to their parallel evolution. Our findings demonstrate that, once integrated into an obligate partnership, newly acquired nutritional symbionts can converge on the long-term evolutionary dynamics of ancient obligate symbionts while undergoing functional specialization.

## Introduction

Microbial symbioses have emerged independently countless times in distant insect clades bringing new functions for their hosts and facilitating the colonization of ecological niches (Moran 2007; Sudakaran, et al. 2017). Colonisation of a nutrient-poor substrate, like blood or phloem-sap, is a clear example where the extended metabolic capacities brought by endosymbionts have enabled insect ecological diversification. Indeed, these diets are poor in essential amino acid and/or vitamins and arthropods that are obligate blood or sap feeders generally rely on obligate endosymbionts to supply them with these essential nutrients (Buchner 1965; Douglas and Prosser 1992; Shigenobu, et al. 2000; Duron and Gottlieb 2020).

Over evolutionary time, hosts and obligate endosymbionts coevolve and become inextricably linked. As a result, many bacterial symbionts of arthropods inhabit specialized host cells called bacteriocytes, usually forming an organ known as the bacteriome (Buchner 1965). This compartmentalization and coevolutionary process have led to convergent modifications of the symbionts. Within these modifications, genomic changes include significantly reduced genome sizes, biased nucleotide compositions, and loss of DNA repair mechanisms when compared to their free-living relatives (Andersson and Andersson 1999; Wernegreen 2002; Toh, et al. 2006; Toft and Andersson 2010). The study of various endosymbiont lineages has allowed a description of the processes leading to this genome reduction and specialisation (Moran and Bennett 2014). These mechanisms include purifying selection acting on genes involved in essential functions for the bacterium and its host. In contrast, genes unnecessary for an intracellular life-style are lost through genetic drift. Vertical transmission with successive bottlenecks in endosymbiont populations also further reduce the effectiveness of selection, even on required genes (Wernegreen and Moran 1999; Moran and Mira 2001; Toft and Andersson 2010). The consequence is a greater rate of fixation of mildly deleterious mutations throughout the symbiont genome. Furthermore, the lack of recombination due to segregation in bacteriocytes and the loss of genes involved in DNA repair/recombination, accelerate the accumulation of irreversible mutations (Moran 1996; Moran and Mira 2001). Within the existing diversity of endosymbiotic systems, the extent of genome reduction and the functional categories of preserved genes vary and depend on the roles carried by the symbiont (McCutcheon and Moran 2012; Luan, et al. 2015; Weglarz, et al. 2018; Choi et al. 2025b).

In some symbiotic systems, the erosion of obligate symbiont genomes has led to the loss of essential metabolic functions, with potentially detrimental effects on host fitness. In such cases, a secondary obligate symbiont that either replaces or complements the primary symbiont metabolic losses is often observed. Instances of such metabolic complementation have been documented across multiple insect groups, including blood feeders such as ticks (Buysse, et al. 2022), and various lineages of sap-feeders within Hemiptera-such as leafhoppers (Takiya, et al. 2006; McCutcheon and Moran 2007), planthoppers (Deng et al., 2023), psyllids (Sloan and Moran 2012; Dittmer, et al. 2023), whiteflies (Santos-Garcia, et al. 2018), scale insects (Husnik and McCutcheon 2016); adelgids (Dial, et al. 2022) and aphids (Manzano-Marín, et al. 2023; Yorimoto, et al. 2025). These systems appear evolutionary unstable, as they are often characterized by high endosymbiont turnover and multiple independent acquisitions throughout host diversification (Koga, et al. 2013; Meseguer, et al. 2017; Matsuura, et al. 2018; Michalik, et al. 2021; Szabó, et al. 2021). However, some dual symbioses deviate from this pattern and have remained remarkably stable over long evolutionary timescales as observed in plant-hoppers (Urban and Cryan 2012; Michalik, et al. 2026) while for many, the evolutionary time span of multi-partner endosymbioses remains poorly documented. In addition, despite hypotheses that functional redundancy following secondary symbiont acquisition may accelerate genome erosion in the primary symbiont (in aphids Lamelas, et al. 2011; Nozaki, et al. 2025; planthoppers Michalik, et al. 2026 and giant scale insects Choi, et al. 2025a), few studies have formally tested this prediction using formal evolutionary rate analyses (but see Vasquez and Bennett 2022; Degnan et al. 2025).

Aphids (Hemiptera: Aphididae) have been intensively studied for their association with their obligate nutritional symbiont, *Buchnera aphidicola* (Enterobacterales: Erwiniaceae) that provides the missing nutrients from their phloem sap diet since their early diversification. These mutualistic partners are localized in bacteriocytes favouring their vertical transmission, this has resulted in long-term cospeciation between aphids and *Buchnera* (Moran, et al. 1993; Clark et al. 2000; Jousselin, et al. 2009). The evolutionary dynamics of the *Buchnera* genome is well documented: it exhibits a reduced genome, a low GC content and a large part of its genetic repertoire is specialised in complementing the host metabolism (Shigenobu, et al. 2000; Tamas, et al. 2002; Chong, et al. 2019). Although aphids were long thought to rely on a single obligate symbiont, recent studies have shown that *Buchnera* is not always the sole provider of essential nutrients. In several aphid lineages, *Buchnera* is complemented by a co-obligate symbiont that contributes to the synthesis of essential B vitamins, most commonly riboflavin (B2) and biotin (B7) (Lamelas et al. 2008; Meseguer et al. 2017; Monnin et al. 2020; Yorimoto et al. 2022; Manzano-Marín et al. 2023). These co-obligate partners belong to diverse Gammaproteobacteria and are thought to have originated from facultative symbionts (Manzano-Marín et al. 2023). The most widespread example is *Serratia symbiotica*, a facultative symbiont that seems to have transitioned to a co-obligate nutritional role in different aphid lineages (Manzano-Marín et al. 2023). However, the long-term stability of these multipartite associations and their evolutionary consequences for both symbiotic partners remain poorly understood.

In the current study, we investigate the evolutionary trajectories of *Serratia symbiotica* and *Buchnera aphidicola* in a clade of aphids that is known to be associated uniquely with this co-obligate duo (Meseguer, et al. 2017). Through comparative genomics, we first analyse the metabolic interdependence between *Serratia* and *Buchnera*. We then use phylogenomics and microscopy to investigate whether obligate *S. symbiotica* derived from a single acquisition event in this lineage and further infer aphid-symbionts co-diversification patterns. Next, we analyse inter-and intraspecific genetic variations in endosymbiont genomes in order to determine whether *Buchnera* and *Serratia* evolve at the same pace. Finally, we investigate genome wide signatures of selection in both endosymbionts in order to assess whether the presence of a co-symbiont significantly changes selective pressures on *Buchnera*. Altogether, these analyses provide an overview of the evolutionary processes that govern the evolution of obligate symbionts in multi-partner endosymbioses.

## Materials and methods

### Aphid collection, DNA extraction and sequencing

We used samples from twelve species from a clade of *Cinara* uniquely associated with *Serratia* (corresponding to Clade A in Meseguer, et al. 2017) and the genome data obtained for *Cinara fornacula* from Meseguer, et al. (2017) (supplementary Table S1). We used aphid colony samples stored in 70% ethanol at 6 °C in the CBGP collection (DOI: 10.15454/D6XAKL). For two aphid species, *Cinara mariana*, *Cinara fornacula,* in order to investigate intraspecific variations in symbiont genomes, we sequenced additional samples from seven and six colonies, respectively (Table S1). For bacterial whole genome sequencing, we used between 15 to 20 aphids from each colony sample and prepared a pooled DNA extract enriched in bacteria as in Jousselin, et al. (2016). Extracted DNA was used to prepare two custom paired-end libraries. Briefly, 5 ng of genomic DNA were sonicated, using the E220 Covaris instrument (Covaris, USA). Fragments were end-repaired, 3’-adenylated, and NEXTflex PCR free barcodes adapters (Bioo Scientific, USA) were added by using NEBNext® Ultra II DNA library prep kit for Illumina (New England Biolabs, USA). Ligation products were then purified by Ampure XP (Beckman Coulter, USA) and DNA fragments (>200 bp) were PCR-amplified (2 PCR reactions, 12 cycles), using Illumina adapter-specific primers and NEBNext ® Ultra II Q5 Master Mix (NEB). After library profile analysis by Agilent 2100 Bioanalyser (Agilent Technologies, USA) and qPCR quantification, using the KAPA Library Quantification Kit for Illumina Libraries (Kapa Biosystems, USA), the libraries were sequenced, using 251 bp paired-end reads chemistry on a HiSeq2500 Illumina sequencer. For the libraries made on *C. mariana* and *C. fornacula* samples to generate intraspecific data, 150 bp paired-end reads chemistry was used.

### Fluorescence in situ hybridization microscopy

We investigated symbiont localisation patterns in individuals from three aphid species: *Cinara palaestinensis*, *Cinara magrhebica*, and *Cinara pinea.* Aphids were first fixed upon collection in Carnoy’s fixative and left overnight, following Koga, et al. (2009). Individuals were then dissected in absolute ethanol, bleached into a 6% solution of H_2_O_2_ diluted in absolute ethanol and then washed with absolute ethanol. Hybridization was performed overnight at 30 °C in standard hybridization buffer and then washed before slide preparation. The embryos of up to 10 individuals were viewed under a TCS SP5 X, Leica confocal microscope. We used two competitive specific probes for *Buchnera* (BLach-FITC [5’-FITC-CCCGTTCGCCGCTCGCCGGCA-FITC-3’] (Manzano-Marín, et al. 2017) and for *Serratia* symbionts in *Cinara* (SeJR5Cinara-Cy3 [5’-Cy3-CCGTYCGCCGCTCGTCACC-Cy3-3’], this study). TIFF-formatted images were exported using the Leica application suite X. Exported images were imported into Inkscape v0.92.4 for building the published figures.

### Genome assembly and annotation

For each library, Illumina reads were right-tail clipped (minimum quality threshold of 20), using FASTX-Toolkit v0.0.14 (http://hannonlab.cshl.edu/fastx_toolkit/). Reads shorter than 75 base pairs (bp) were dropped. Additionally, PRINSEQ v0.20.4 (Schmieder and Edwards 2011) was used to remove reads containing undefined nucleotides as well as those left without a pair after the filtering and clipping process. The resulting reads were assembled, using SPAdes v3.10.1 (Bankevich, et al. 2012), with the option --only-assembler option and k-mer sizes of 33, 55, 77, 99 and 127. From the resulting contigs, those that were shorter than 200 bp were dropped. The remaining contigs were binned, using results from a BLASTX (Altschul, et al. 1997) search (best hit per contig) against a database consisting of the pea aphid’s proteome and a selection of aphid’s symbiotic bacteria previously found in *Cinara* species and free-living relatives’ proteomes. The assigned contigs were manually screened using the BLASTX web server (*vs.* the nr database) to insure correct assignment.

We generally obtained for each sample a contig of about 16 kb that corresponded to a complete aphid host mitochondrial genome. Those were annotated using Mitos WebServer (Bernt, et al. 2013) followed by manual curation in Geneious v11.1.5. The cox1 genes retrieved in our assemblies were systematically used in BlastN searches against our own curated barcode database (based on specimen vouchers) (Coeur d’acier, et al. 2014) to confirm species identification and ensure that we did not have any contamination. We also obtained a complete *Buchner*a chromosome generally into two contigs and at the most 40 (for *Cinara pruinosa*) and *Serratia* chromosomes were scattered into 2 to 13 contigs (Table S2). The contigs were ordered using Mauve Contig Mover algorithm implemented in Geneious v11.1.5 (https://www.geneious.com) using as references *Serratia symbiotica* and *Buchnera aphidicola* associated with *C. fornacula* studied by Meseguer, et al. (2017). For the 12 newly sequenced aphid species, and the previously sequenced endosymbionts of *C. fornacula*, the resulting *Buchnera* and *Serratia* chromosomes were annotated by submitting the data to the MicroScope platform pipeline (Vallenet, et al. 2019). This annotation was followed by manual curation of the coding sequences (CDS) on Geneious v11.1.5 and the MicroScope platform using BlastX against reference sequences of *Buchnera aphidicola* and *Serratia symbiotica*. The resulting CDSs were considered to be putatively functional if all essential domains for the function were found, if a literature search supported the truncated version of the protein as functional in a related organism, or if the CDSs displayed truncations but retained identifiable domains (as shown by Tamas, et al. 2008).

### Intraspecific variation in symbiont genomes in two Cinara species

Illumina reads were processed as previously described to obtain *Serratia* and *Buchnera* chromosome assemblies for each sample. When several contigs were obtained for a symbiont genome, scaffolding was conducted by whole genome alignment between resulting contigs and reference endosymbiont genomes of *C. mariana* and C. *fornacul*a obtained above using Mauve Contig Mover algorithm implemented in Geneious v11.1.5. Finally, genome pair-wise alignments were visually verified and annotations were transferred from reference genome using Geneious tools (“transfer annotation”). Manual curation of start and stop codons of all CDSs was conducted in Geneious v11.1.5.

### Phylogenetic analyses

#### Cinara mitochondrial phylogeny

In order to reconstruct the phylogeny of the aphid hosts (clade A of *Cinara* species) we used all thirteen mitochondrial CDSs from the 13 newly sequenced *Cinara* species. We also selected mitochondrial genomes from three outgroup species (*Acyrtosiphon pisum, Cinara cedri and Cinara tujafilina*) retrieved from NCBI (Table S3).

For each gene, DNA sequence alignments were conducted using Macse v2.03 (Ranwez, et al. 2018). We then removed divergent and ambiguously aligned blocks using Gblocks v0.91b (Talavera and Castresana 2007) and concatenated the resulting alignments in one matrix. A partitioned scheme was set with one data block defined for each codon position in each gene. Substitution models on the concatenated datasets was selected using ModelFinder (Kalyaanamoorthy, et al. 2017) (MF option implemented in IQ-TREE 1.6.8) using the Bayesian Information Criterion and ML phylogenetic inferences were conducted using IQ-TREE 1.6.8 (Nguyen, et al. 2014) with best model and 1000 ultrafast bootstrap. We also ran a Bayesian phylogenetic inference using MrBayes v3.2.7 (Ronquist, et al. 2012) with the best PartitionFinder scheme available in MrBayes and running two independent analyses with four chains each for 3,000,000 generations and checked for convergence.

#### Buchnera and Serratia phylogenies

For phylogenetic inference, orthologous protein-coding sequences were identified separately for each symbiont dataset using Orthofinder v2.5.4 (Emms and Kelly 2019). The *Buchnera* dataset comprised the 13 genomes generated in this study together with four genomes from outgroup species, whereas the *Serratia* dataset included the 13 genomes generated in this study and three outgroup strains, including *Serratia marcescens* and facultative *Serratia symbiotica* strains associated with aphids (Table S3). We respectively obtained 357 genes shared by all *Buchner*a s and 546 genes shared by all *Serratia*. For each symbiont species lineage, each genes were aligned as nucleic acid sequences using MAFFT v7.450 (maxiterate 1,000 localpair) (Katoh and Standley 2013). Divergent and ambiguously aligned blocks were removed using Gblocks, genes were then concatenated. Substitution models on the concatenated datasets for each symbiont were selected using ModelFinder (MF option implemented in IQ-TREE 1.6.8) using the Bayesian Information Criterion. ML phylogenetic inferences were conducted using IQ-TREE 1.6.8 (Nguyen, et al. 2014) with best model and 1000 ultrafast bootstrap. Bayesian phylogenetic reconstructions were also conducted with MrBayes v3.2.7 MPI under the GTR+I+G model, running two independent analyses with four chains each for 300,000 generations (sampling every 1000 generations and 20% discarded as burn-in).

### Examining patterns and pace of genome evolution in co-evolving symbionts

#### Estimating evolutionary rates in Serratia and Buchnera

Calibration of *Cinara*’s phylogeny was conducted in Beast v.1.10.4 (Drummond and Suchard 2010) over the previously described mitochondrial DNA matrices (excluding outgroups) using the best partition scheme and models found previously. The selected priors were a Birth-Death model with an uncorrelated log-normal molecular clock and fixed tree topology using the ML tree obtained as previously described. We used previous phylogenetic inquiries of *Cinara* to define three calibration points for the estimation of substitution rates: the crown age of *Cinara* clade A was dated as 25 My (Million years) by Meseguer, et al. (2015). Tree root was thus calibrated here using a normal prior distribution: mean of 25, standard deviation of 3. We used the two oldest nodes of the tree as others calibration points (Figure 2, node X: mean of 21, stdev of 2 and node T: mean of 21.7, stdev of 2). We ran two MCMC chains of 150 million generations in Beast and assessed Convergence using Tracer 1.7 (Rambaut, et al. 2018). After removing 20% of the samples as burn-in, we used TreeAnnotator 1.8 to generate a maximum clade credibility tree with the median age estimate of nodes and average substitution rates. As there was a perfect congruence between the phylogenies of *Serratia*, *Buchnera* and their *Cinara* hosts, we used aphid host node dates to calibrate symbiont phylogenies. Substitution rates in both symbiont phylogenies were estimated with Beast using settings previously described for the aphid mitochondrial data, on both symbiont datasets, using a GTR+I+G4 substitution model, and two MCMC chains of 300 million generations.

To compare the pace of genome evolution in *Serratia* and *Buchnera*, we extracted root-to-tip branch lengths (substitutions per site) from maximum-likelihood (ML) trees inferred for Clade A in each symbiont (undated trees), following the approach of Arab, et al. (2020). Pearson correlation analyses were performed in R (R Core Team 2022) to assess correlation in branch lengths between corresponding lineages of the *Serratia* and *Buchnera* phylogenies, as well as between each symbiont and the aphid mitochondrial phylogeny. To test comparability based on homologous gene sets, we repeated the analyses using ML trees reconstructed from genes shared between *Serratia* and *Buchnera* (i.e. CDS with the same annotation in both symbionts). For each symbiont, DNA matrices were restricted to orthologous genes present in both lineages, and phylogenies were inferred as described above. Root-to-tip branch lengths were then re-estimated and the correlation analyses repeated.

#### Gene loss rates in Serratia

As *Serratia* genomes were characterized by variations in CDS contents (pseudogenes or missing genes), from a matrix of absence/presence of functional CDS in *Serratia* genomes, we computed the number of CDS losses events on *Serratia* calibrated phylogeny obtained above using the software Count (Csűrös 2010) and assuming a CDS cannot be gained once lost. For each branch, gene loss rates were then calculated by dividing the number of losses assigned to a branch by its temporal duration, and mapped onto the time-calibrated *Serratia* phylogeny using Ape (the R statistical software R Core Team, 2022).

#### Test of selection

Using the 357 coding sequences (CDSs) shared among all 13 *Buchnera* genomes, the 545 CDSs shared among all 13 *Serratia* clade A genomes, and the maximum-likelihood phylogenies inferred above, we estimated the ratio of nonsynonymous to synonymous substitution rates (dN/dS, ω) and tested for signatures of selection on each orthologous gene. Estimates of ω were obtained using the CODEML program implemented in PAML v4.9 (Yang, 2007). Under this framework, ω > 1 indicates positive selection, ω = 1 is consistent with neutral evolution, whereas 0 < ω < 1 indicates purifying selection, with lower values reflecting stronger selective constraint. Because vertically transmitted endosymbionts are expected to evolve predominantly under purifying selection, variation in ω was interpreted primarily as reflecting differences in the strength of selective constraint rather than adaptive evolution. To identify if some genes evolved under positive selection, we applied two complementary approaches. First, we used the branch-site unrestricted statistical test for episodic diversification (BUSTED) implemented in HyPhy (Murrell et al., 2015; Kosakovsky Pond et al., 2020), which tests whether a gene has experienced episodic positive selection affecting at least one site on at least one branch of the phylogeny. Second, we performed the site-model likelihood ratio tests M1a versus M2a and M7 versus M8 implemented in CODEML (Anisimova et al., 2001), which compare models that restrict ω to values ≤ 1 with models allowing a class of sites to evolve with ω > 1. Statistical significance was assessed using likelihood ratio tests, with the significance of nested models evaluated by chi-squared tests.

To test whether the co-occurrence of symbionts with gene redundancy, affected selective constraints on shared CDS, potentially further accelerating genomic erosion in symbionts, we analysed variations in dN/dS values. Under the hypothesis that redundancy in metabolic function facilitate the decay of the genes coding for the same function, we expect a significant impact of the shared status of CDS on their dN/dS values, when they are involved in symbiotic functions (host metabolism). We used a generalised linear model (GLM) with log-transformed dN/dS as a response variable and Gaussian error distribution. We specified three explanatory variables: strain (with two categories: *Buchnera* N=355 and *Serratia* N=545); CDS functions (those were reduced to five categories derived from our annotation: Expression, replication and genome maintenance N=370; Cell processes and signalling N=203; Host provisioning N=143; Other metabolism N=162; poorly characterized = 22) and status (with two categories: shared N=496 or not-shared N=404). All factors and their interactions were tested. The goodness of fit of the model was assessed by examining difference of deviance with or without the factor tested. When a factor showed a significant effect, we conducted post hoc analyses using estimated marginal means implemented in the *emmeans* package. Pairwise comparisons between levels of factors were performed using Tukey-adjusted contrasts. In particular, we tested differences between shared and non-shared CDS within each functional category. We then restricted the dataset to genes shared between *Buchnera* and *Serratia*, plotted the dN/dS values of orthologous genes, and tested the correlation between the two symbionts separately for each functional category using Pearson correlation. All statistical analyses were performed using the R statistical software (R Core Team, 2022).

#### Coevolution and pace of genome evolution at the intraspecific level

For both species for which we gathered intraspecific data (*C. mariana* and *C. fornacula*), we reconstructed *Serratia* and *Buchnera* phylogenies to examine patterns of phylogenetic congruence at the intraspecific level. Maximum likelihood phylogenetic analyses were conducted as described for the clade A phylogeny. For each aphid species and for each symbiont lineage, we extracted all coding genes from annotated *Serratia* and *Buchnera* chromosomes using Geneious v11.1.5 and kept orthologous genes. Each ortholog gene was then aligned using MAFFT v7.450 (--maxiterate 1000 --localpair). Divergent and ambiguously aligned blocks were removed using Gblocks v0.91b and gene alignments were concatenated. Phylogenetic reconstruction was conducted with IQ-TREE v1.6.8 (1000 ultrafast bootstraps) using the GTR+F+G4 model for both symbionts as selected by –MF in IQ-Tree.

For each aphid species and each endosymbiont strain, we conducted whole genome alignment of annotated genomes in Geneious v11.1.5 using Mauve, we then extracted tables of polymorphism using Geneious function “Find variations/SNP” on the resulting alignments. We then inferred deletion/insertion and single nucleotide polymorphism (SNP) numbers and ratios (per kb) in *Serratia* and *Buchnera* datasets for both *C. mariana* and *C. fornacula*.

## Results

### Endosymbiont genome characteristics and metabolic complementarity

The sequences of the chromosomes of thirteen *Buchnera-Serratia* pairs from *Cinara* species were successfully sequenced, assembled and annotated. *Buchnera* chromosomes are small (from 451 to 464,5 Kbp) and G+C poor (24,7 to 28,1%) (Supplementary Table S1, Figure S1), their gene contents are highly conserved (365 to 372 CDS). As for *Serratia* symbionts, once scaffolded we retrieved a nearly complete bacterial chromosome for all strains; genomes have a size between 1.10 and 1.28 Mb with an average G+C content between 28.3 and 35.8 % (Supplementary Table S2 and Figure S2). Their gene contents exhibit more variations than *Buchnera* (631 to 681 protein coding genes). Finally, a single conserved copy of 16S/23S/5S rRNA was found in all 13 *Buchnera* genomes and 31 and 33 to 35 tRNAs were found in *Buchnera* and *Serratia* respectively.

To infer the dependence of both *Buchnera* and *Serratia* for nutritional complementation of the aphid, we compared the gene repertoire related to the biosynthesis of EAAs, vitamins, and co-factors in the 13 pairs of endosymbiont genomes and looked at overlap and complementarity in these nutritional functions (Figure 1). *Serratia* has lost most EAAs’ biosynthetic genes, and therefore depends on *Buchnera* for the provision of these nutrients. Similarly, to other co-obligate symbionts of Lachninae studied so far, *Serratia* strains encode genes that are missing from *Buchnera* to accomplish the biosynthesis of both riboflavin (full pathway) and biotin (*bioA*, *bioD* and *bioB* genes).

**Figure 1:**
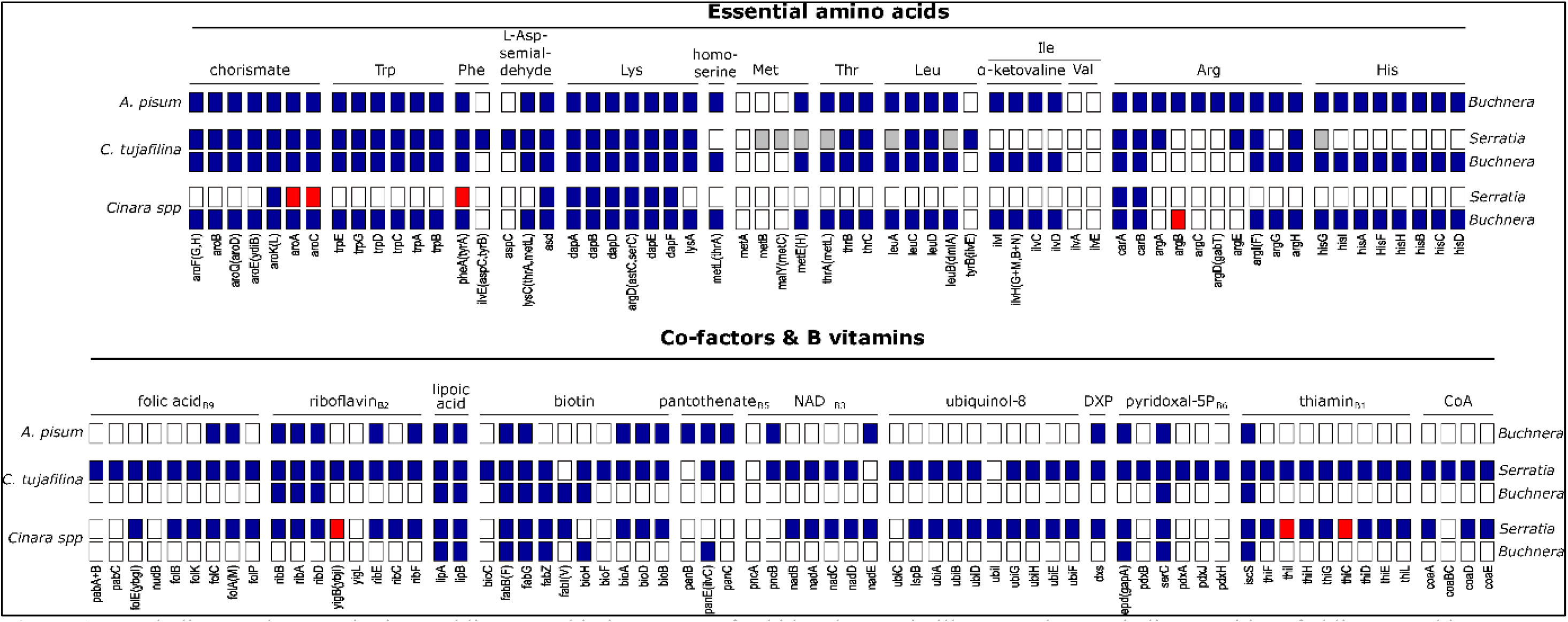
Metabolic complementarity in co-obligate symbiotic systems of aphids. The matrix illustrates the metabolic capacities of obligate symbionts across different aphid species. “*Cinara* spp.” refers to *Cinara* species newly sequenced in this study. The genome of *Buchnera* from the pea aphid (*Acyrtosiphon pisum*), which harbors *Buchnera* as its sole nutritional symbiont, is shown for comparison, as well as the symbiont pair associated with *Cinara tujafilina*, which hosts a larger *S. symbiotica*. Blue boxes indicate the presence of genes, white boxes indicate their absence, red boxes represent genes lost in at least one *Cinara* species, and grey boxes indicate pseudogenes. At the top of each column, the enzyme or pathway catalyzing the reaction is indicated, while at the bottom, the compound produced by the corresponding enzymatic step is shown. Symbiont taxon names are given to the right of each row, and host taxon names to the left.

**Figure 2:**
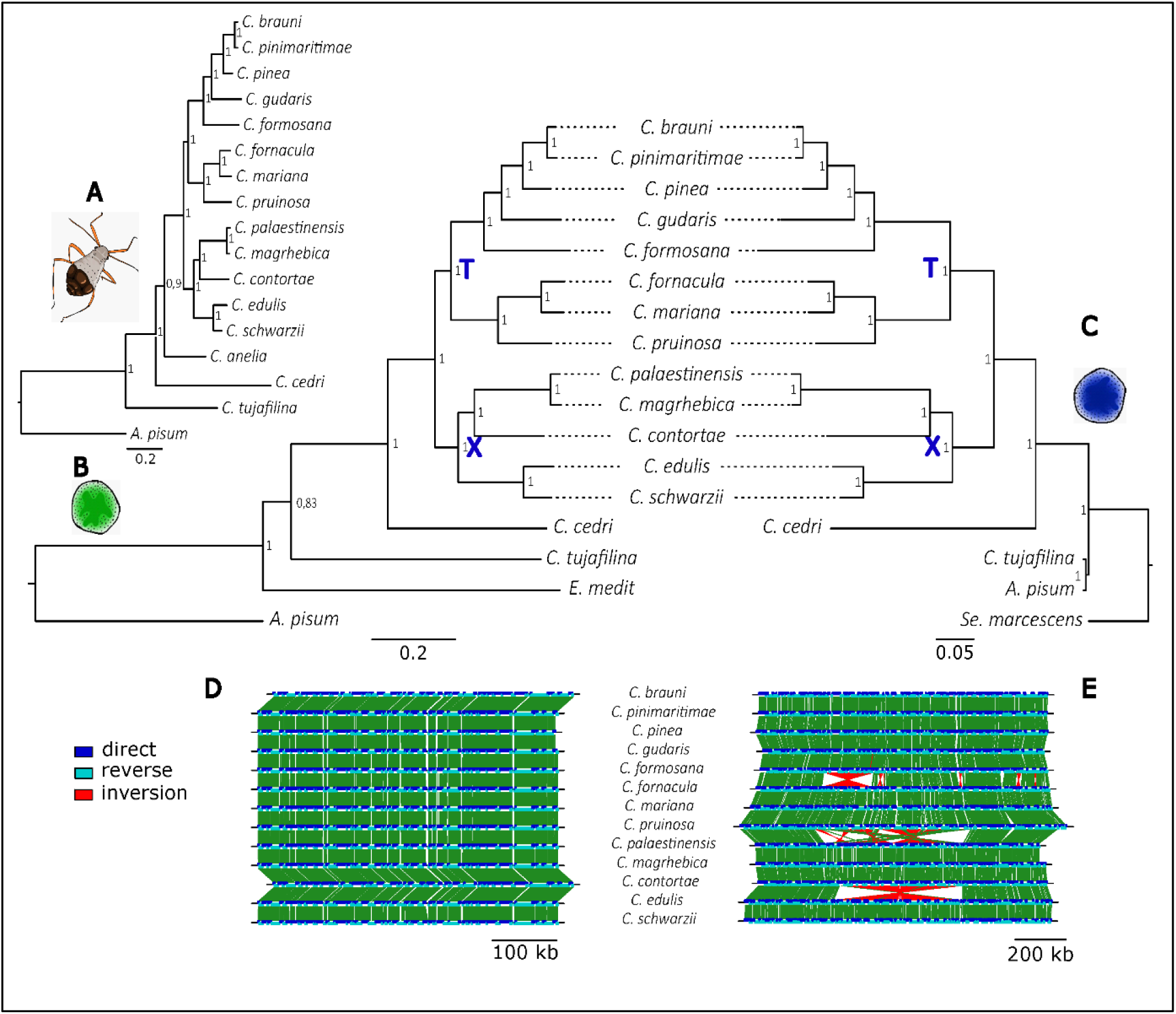
Phylogenetic reconstructions inferred with Bayesian analyses of *Buchnera*, *Serratia* chromosomes and host mitochondrial genomes. **A** Phylogenetic reconstruction of *Cinara* species of clade A, using aphid mitogenomes (*C. tujafilina* and *A. pisum* are placed as outgroups). **B** Phylogeny of the associated *Buchnera* strains (*Buchnera* associated with *C. tujafilina*, *A. pisum* and *E. mediterraneus* are used as outgroups). **C** Phylogenetic reconstructions of associated *Serratia* srains, (obligate *S.symbiotica* from *C. tujafilina*, facultative *S. symbiotica* from A*. pisum* and *Serratia marcescens* are used as outgroups). Numbers at each node indicate posterior probabilities. The nodes used for time calibration are represented by X and T. **D** Gene synteny of the thirteen *Buchnera* chromosomes. **E** Gene synteny of the thirteen *Serratia* chromosomes.

In addition, *Serratia* encode for proteins involved in the biosynthesis of thiamine (*thiF*, *thiS*, *thiI*, *thiH*, *thiG*, *thiC*, *thiD*, *thiE* and *thiL*), as the co-obligate symbiont *E. haradaeae* of another clade of *Cinara* (Manzano-Marín, et al. 2020). *Buchnera* genomes are highly conserved and exhibit nearly perfect synteny (Figure 2D and S1). In *Buchnera* genomes, CDS content variation was limited to 11 CDS (*trmD*, *murE*, *hinT*, *argB*, *fliO*, *flgI*, *fis*, *yciC*, *FlgH*, *yoaE*, *orn*; Figure S1) each of which was lost in at least one lineage. In contrast, *Serratia* genomes displayed several inversions and greater CDS content variability (Figures 2C and S2, Table S4). The observed inversion positions were systematically found within contigs, those were all checked manually.

### Phylogenetic history of Cinara di-symbiotic system

For each dataset, ML and Bayesian phylogenies were highly congruent and well resolved (Figure 2 and Supplementary-material). Phylogenetic analyses indicate that all *Serratia symbiotica* of the *Cinara* species used in this study form a well-supported monophyletic group (Figure 2, Figure S2). Moreover, the comparison between *Cinara* mitochondrial genomes*, Buchnera* and *Serratia* phylogenies suggests a perfect congruence of their evolutionary histories (Figure 2).

### Endosymbiont genome evolution

Beast analyses inferred substitution rates of 10.5*10^-^9 substitution/sites/years (s/s/y) for *Buchnera* and 9.25*10^-^9 s/s/y for *Serratia*. For the mitochondrial genomes, a rate of 4.66*10^-^ 9 s/s/y was obtained.

#### Evolutionary rate correlation

Comparison of parallel branches in host and symbiont ML phylogenies (Figure 3A) revealed a significant correlation between the substitution rates in mitochondrial and bacterial genomes (host-*Buchnera*: R = 0.82, *P* = 7.7^e^-10; host-*Serratia:* R =0.62, *P* = 3.3^e-6^; Figure 3B). Significant correlation was also found between the substitution rates of *Serratia* and *Buchnera* (R= 0.87, *P* = 1.5^e-11^; Figure 3C). The latter remained valid when only genes shared by both symbionts were used in the phylogenetic reconstruction (R= 0.88, *P* = 2^e-11^, Figure S3).

**Figure 3:**
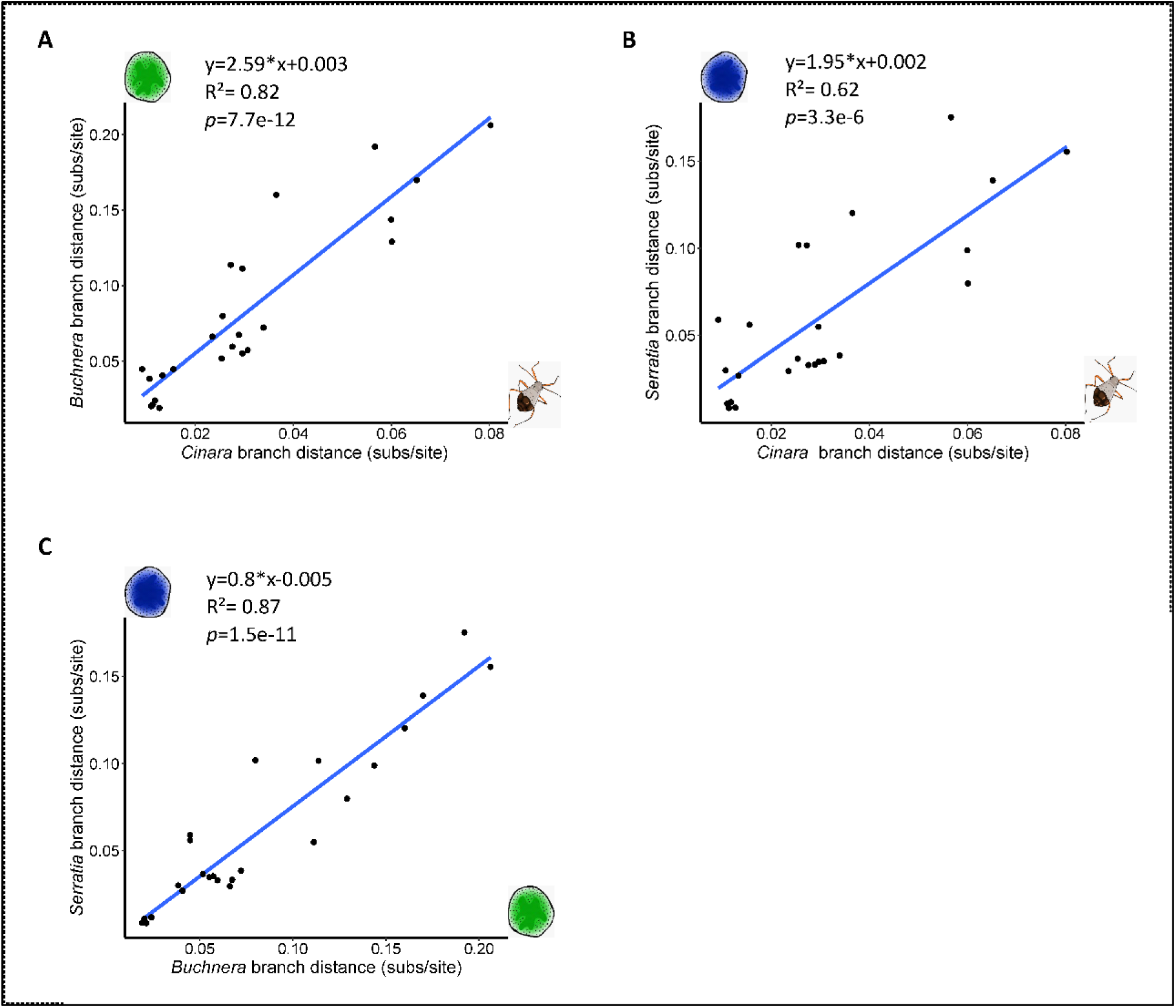
Comparison of evolutionary rates of *Serratia* and *Buchnera* and the associated *Cinara* mitochondrial genomes. A) Correlation of root-to-tip branch lengths (in subs/site) from maximum-likelihood phylogenies of aphids and *Buchnera.* B) Correlation of root-to-lip branch lengths (in subs/site) from maximum-likelihood phylogenies of aphids and *Serratia.* C) Correlation of root-to-tip branch lengths (in subs/site) from maximum-likelihood phylogenies of *Buchnera* and *Serratia.* For each graph, the R^2^ value, p-value, and regression line are shown.

#### Rate of gene losses in Serratia

The mapping of number of CDSs lost onto the *Serratia* calibrated phylogeny showed that gene loss rates varied across the branches. In total, 145 CDS loss events were inferred onto the phylogeny and some CDS were lost up to five times (Figure 4, Table S4). Gene losses appeared in all gene categories (Figure 4, Table S4). Estimated rates were heterogeneous across the phylogeny, they ranged from 0 to 32 CDS losses per Myr (mean = 3.84; median = 1.90). Most recent branches exhibiting relatively low rates, while a small number of branches showed markedly elevated rates (Supplementary Table S4, Figure S4).

**Figure 4:**
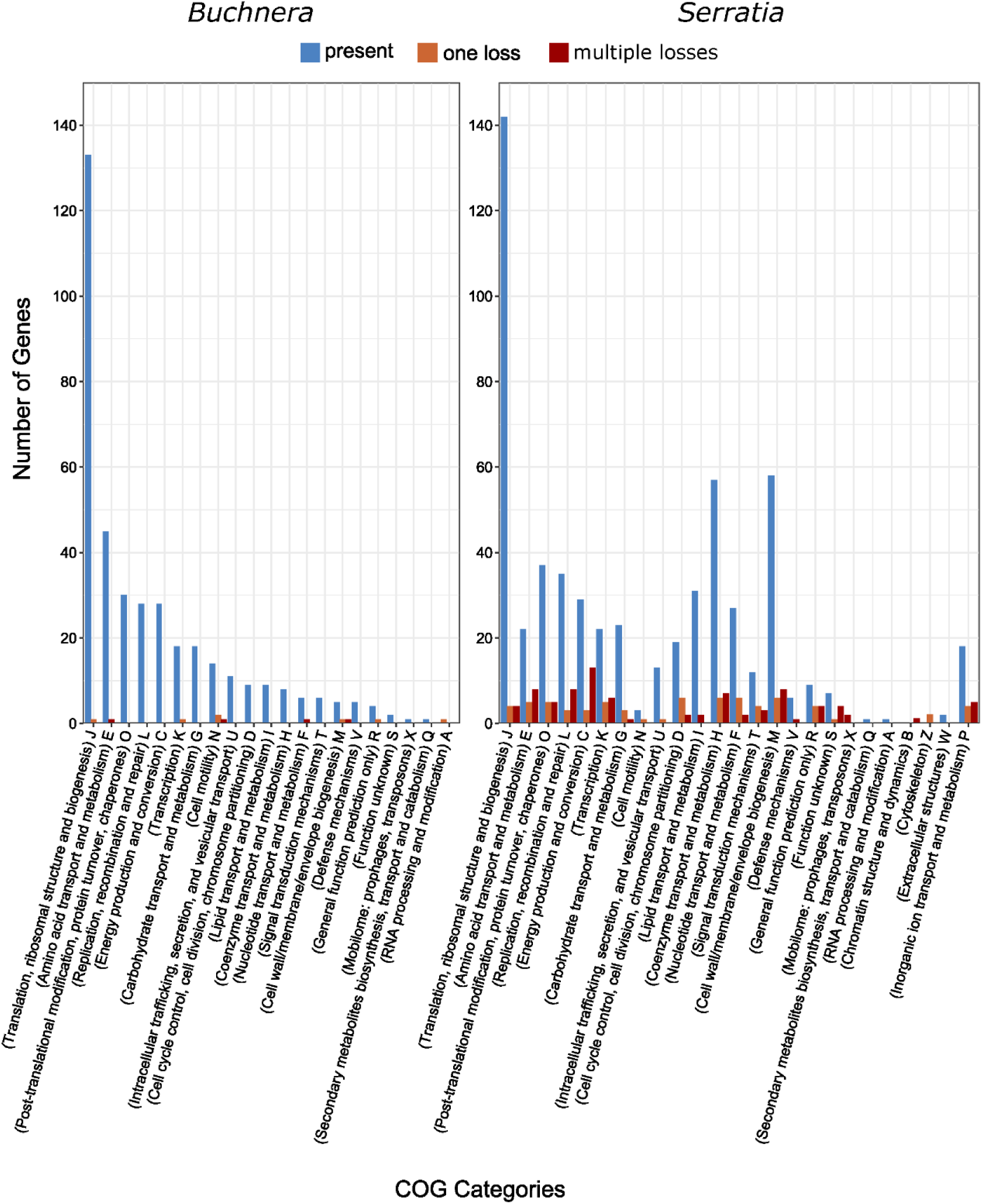
Bar chart showing the number of genes lost and present across *Buchnera* (right) and *Serratia* (left) chromosomes from *Cinara* species. Genes are binned into their Clusters of Orthologous Groups (COGs) functional categories (Tatusov, et al. 2003). Bars are color coded according to the number of independent losses (i.e., genes lost in none of the species in blue, genes lost once in orange and genes with multiple independent losses in red).

#### Signature of selection across endosymbiont genomes

Using M0 model in Codeml, we found mostly genes undergoing strong purifying selection in *Buchnera* with very low dN/dS values (avg. ω = 0.0625 [range = 0.0053-0.0999, n = 276]), few genes undergoing more relaxed purifying selection (avg. ω = 0.1328 [range = 0.1003-0.2556, n = 79]) and no gene under positive selection (Table S5). The M1a-M2a and M7-M8 codeml nested models identified 1 to 51 genes under positive selection, respectively, while Busted analysis revealed 10 genes under positive selection (Table S5). However, none of the genes displayed strong evidence of positive selection supported by all three tests.

Using M0 model in Codeml, we found that in *Serratia symbiotica* many genes were undergoing strong purifying selection (avg. ω = 0.0682 [range = 0.0001-0.0999, n = 135]), but more genes were experiencing relaxed purifying selection (avg. ω = 0.1590 [range = 0.1000-0.4025, n =411]) (Table S6). As for *Buchnera*, no gene exhibited robust evidence of positive selection supported by all three tests. Indeed, we detected 3 to 48 genes with signals of positive selection using the M1a-M2a and M7-M8 Codeml nested models, respectively, while Busted analysis identified 9 genes under positive selection (Table S6).

Using GLM models, we found a significant effect of the interaction term status* function (Table 1), meaning that depending on gene function, the fact they were shared or not between symbionts, affected dN/dS values. Post-hoc contrasts revealed that non-shared genes exhibited significantly higher dN/dS values than shared genes in the *cellular processes and signaling* category, whereas no significant effect of status was detected for other functional categories (all *p* > 0.45) (Table 1, Figure S5). In addition, we found an overall correlation of dN/dS values of shared genes in all functions except for host-provisioning genes, but this gene category includes only 16 genes (Figure S6).

**Table 1:** Results of the GLM analysis of the dN/dS values calculated wih Codml, function refers to COG functions simplified into four categories, strain refers to endosymbiont identity (*Buchnera/ Serratia*), status to whether the gene was found in both symbionts (shared) or not (not shared). Contrasts correspond to pairwise comparisons of status within each functional category (emmeans).

| Model term removed | df | Deviance | F | p |
| --- | --- | --- | --- | --- |
| none |  | 277.34 |  |  |
| status(shared/non-shared) × strain × function | 3 | 277.49 | 0.1599 | 0.9 |
| status × strain | 1 | 277.68 | 0.6023 | 0.47 |
| strain × gene function | 4 | 279.06 | 1.2532 | 0.28 |
| **status x gene function** | **3** | **283.63** | **6.5261** | **0.0002\*\*** |
| **strain** | **1** | **337.33** | **185** | **<0.000 \*\*\*** |
| Gene functional category | Estimate | SE | t | P |
| **Cellular processes and signaling** | **0.428** | **0.084** | **5.07** | **<0.0001** |
| Expression, replication, genome maintenance | -0.014 | 0.082 | -0.17 | 0.87 |
| Host provisioning | 0.076 | 0.103 | 0.74 | 0.46 |
| Metabolism | -0.047 | 0.097 | -0.48 | 0.63 |

We found a perfect congruence between *Buchnera* and *Serratia* phylogenies within each *Cinara* species confirming parallel history of symbionts at this fine evolutionary scale (Figures S7 and S8). We then analysed polymorphisms and indels at the intraspecific level (Table 2). In both species, we found strikingly similar patterns of mutations between co-existing symbionts. In the two species, *Buchnera* and *Serratia* exhibit similar number of SNP/kb in coding and non-coding sequences and in both cases non-coding regions exhibited higher (almost twice as high) numbers of substitutions. Numbers of indels are 50 to 100 times higher in non-coding regions than coding regions. In *C. mariana*, higher numbers of indels/ kb were found in *Serratia* than *Buchnera*.

**Table 2:** Mutational patterns in genomes of *Buchnera* and *Serratia* in *Cinara mariana* and *Cinara fornacula*. na: not applicable, Ncoding: non coding, indels: insertion-deletion.

| Aphid species | Symbiont | DNA partition | length (bp) | nb. SNPs | dN/dS | SNPs/kb | total indels | indels/kb | SNP/indels |
| --- | --- | --- | --- | --- | --- | --- | --- | --- | --- |
| *C. mariana*<br>(7 genomes) | *Buchnera* | coding | 369243 | 36320 | 0.33 | 98.36 | 89 | 0.24 | 408.09 |
|  |  | Ncoding | 78838 | 11374 | na | 144.27 | 1146 | 14.54 | 9.92 |
|  | *Serratia* | coding | 626723 | 44554 | 0.34 | 71.09 | 88 | 0.14 | 506.3 |
|  |  | Ncoding | 556968 | 84760 | na | 152.18 | 24041 | 43.16 | 3.53 |
| *C. fornacula*<br>(7 genomes) | *Buchnera* | coding | 370866 | 8587 | 0.35 | 23.21 | 23 | 0.06 | 373.35 |
|  |  | Ncoding | 76807 | 3465 | na | 46.82 | 486 | 6.39 | 7.13 |
|  | *Serratia* | coding | 636090 | 10368 | 0.38 | 16.30 | 58 | 0.09 | 178.76 |
|  |  | Ncoding | 524728 | 20595 | na | 39.30 | 3408 | 6.50 | 6.04 |

### Localisation of Serratia symbiotica in Cinara species

FISH in three *Cinara* species revealed that *Serratia* were exclusively present inside bacteriocytes that were separated from *Buchnera* bacteriocytes, but distributed within the aphid body area known as the bacteriome (Figure 5). *Serratia* also present coccoid cell shape smaller than *Buchnera* cells as previously described for other endosymbionts from other *Cinara* species with highly reduced symbiont genomes (Manzano-Marín, et al. 2017; Manzano-Marín, et al. 2020).

**Figure 5:**
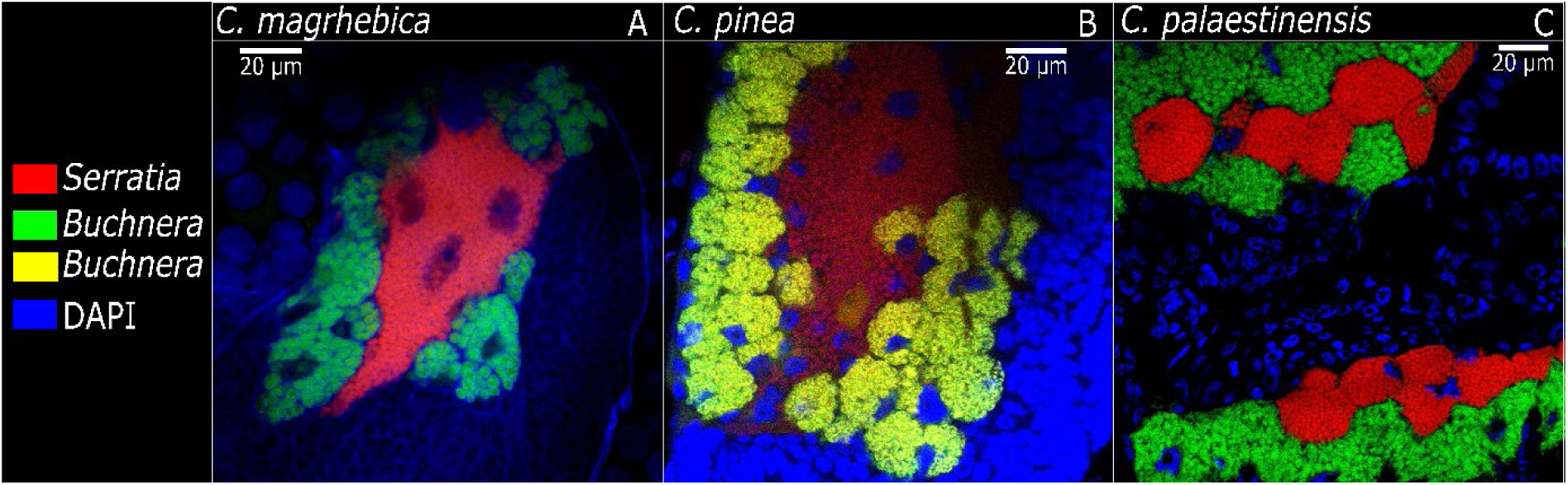
Merged FISH microscopic images of aphid embryos from selected *Cinara* aphids. *Serratia* signal is shown in red, *Buchnera* signal is in green or yellow on figure B, and DAPI’s (staining DNA, highlighting host nuclei) in blue. The name of each species is shown at the top of each panel.

## Discussion

### Serratia symbiotica has co-speciated with a clade of aphids as a co-obligate nutritional partner

We confirmed that a strain of *Serratia symbiotica* is capable of complementing *Buchnera* for its lost symbiotic functions (i.e. the biosynthesis of B vitamins Figure 1) in a clade of *Cinara* aphids and exhibits the typical signature of a long-term obligate association with its hosts, *i.e.* a small genome of ∼1 Mbp and low G-C contents throughout its diversification (Figure S1). Phylogenomic analyses revealed parallel phylogenetic histories of the three partners of this mutualism supporting a scenario of an ancestral acquisition of *Serratia* followed by cospeciation with *Buchnera* and associated aphids for about 25 My (Meseguer, et al. 2015) (Figure 2). This parallel history of symbionts is also observed at the intraspeific level in two aphid species and altogether our results support strict vertical transmission of an obligate *Serratia* in this clade of *Cinara*. These inferences are further corroborated by the presence of *Serratia* specific bacteriocytes in the aphid bacteriome that are very similar in shape, size and localization across three *Cinara* species (Figure 5). Their proximity and similarity to *Buchnera* bacteriocytes raise the possibility for the co-symbiont to follow the paths of vertical transmission of the primary symbiont (Koga, et al. 2012). This coevolutionary history contrasts with the instability of the di-symbiotic systems observed in several clades of Hemipterans where co-symbiont replacements have been observed at different evolutionary time-scales (Rosenblueth, et al. 2012; Koga, et al. 2013; Husnik and McCutcheon 2016; Dial, et al. 2022; Zhang, et al. 2023; Choi et al. 2025b). Our results mirrors those of Manzano-Marín (2020) that showed long-term cospeciation of *Erwinia* and *Buchnera* in another clade of *Cinara* aphids. This indicates that stable associations between *Buchnera* and a new co-obligate symbiont has evolved multiple times in aphids, with different bacterial lineages. For the first time, we show the possibility of a long-term co-diversification pattern of *Serratia symbiotica* with *Buchnera* and their aphid hosts.

In addition to compensating for *Buchnera*’s lost genes, the aphid-associated *Serratia* strains in our study retain a thiamine-biosynthetic capability, a function that is rarely observed in *Buchnera* (Manzano-Marín, et al. 2023). This points towards the possibility that the acquisition and maintenance of *Serratia* as a nutritional symbiont of *Cinara*, is associated with the advantages conferred by this newly recruited function. This hypothesis was put forward for *Erwinia*-associated *Cinara* species that also hold a thiamine biosynthesis capacity (Manzano-Marín, et al. 2020). It is possible that the conifer hosts (*Picea* Piceoideae, *Larix* Laricoideae, but also *Pinus Pinoideae*) on which these aphids live have low thiamine levels, and that a symbiont capable of synthetizing this vitamin is selected and maintained in aphid populations feeding on certain conifers over evolutionary times. This hypothesis is reinforced by the fact that the only *Buchnera* strains retaining this thiamine synthesis capability, not associated to a co-obligate partner, are the *Buchnera* from two aphid genera, *Neophyllaphis* and *Mindarus*, that also feed on conifers, namely Araucariales and firs (*Abies spp. Pinales: Pinaceae: Abietoideae)* (Manzano-Marín, et al. 2023). Comparative data on thiamine content of the sap of different aphid plants will be needed to validate our hypothesis.

Alternatively, the stability of this symbiotic system may also stem from a relative rarity of alternative symbionts that could rescue *Buchnera* in the aphid clade studied. Once *Buchnera* has irreversibly lost metabolic genes, it opens a niche for a symbiont to fulfil the lost symbiotic functions and maintain the systems. Symbiont replacements in such multi-partner systems may therefore occur through successive substitutions among symbiont lineages that are functionally equivalent from the host’s perspective and compete for the same niche. Such a process has been proposed for *Cinara* aphids (Meseguer et al. 2017) but also in the dynamic dual endosymbioses of *Amblyomma* ticks (Buysse et al. 2022). Under this scenario, the absence of suitable competitors could explain why some symbiotic associations remain stable over long evolutionary timescales whereas others experience recurrent symbiont turnover. Comparative data on facultative symbiont diversity in aphid lineages will be needed to test this hypothesis.

In any case, given the diversity of co-obligate symbionts found across aphids (Manzano-Marín et al. 2023; Yorimoto et al. 2025), and particularly within Lachninae (Manzano-Marín et al. 2017; Meseguer et al. 2017), the strict co-speciation pattern observed here is unlikely to be representative of all multipartite endosymbiotic systems of aphids. This study system provides the opportunity to further investigate the factors triggering symbiont replacements.

*Cospeciating* Serratia *and* Buchnera *evolve at the same pace and exhibit similar patterns of molecular evolution*

Using a Bayesian phylogenetic framework that models rate heterogeneity across the tree and calibration uncertainty, we found an average substitution rate of 10.5*10^-9^ (s/s/y) for *Buchnera*. Our estimates place the long-term substitution rate of *Buchnera* within the range reported in previous studies (Brynnel et al. 1998; Clark et al. 2000; Tamas et al. 2002; Gomez-Valero et al. 2007; Perez-Brocal et al. 2011) (Table 3), reinforcing the view that rates of molecular evolution in this primary endosymbiont have remained relatively stable across aphid lineages. More importantly, our analyses reveal that *Serratia* evolves at a nearly identical rate (i.e. 9.25*10^-9^ s/s/y) despite its much more recent integration into the symbiosis. The strong correspondence of branch lengths between the two symbiont phylogenies, including analyses restricted to orthologous genes shared by both partners, indicates that their evolutionary dynamics have become tightly coupled over millions of years (Figure S3). Such rate coupling suggests that the two symbionts are exposed to similar evolutionary constraints. Because substitution rates in bacterial endosymbionts are influenced by generation time, effective population size, and the strength of genetic drift (Degnan et al. 2005), the parallel evolutionary rates observed here imply that *Buchnera* and *Serratia* experience comparable demographic and ecological conditions within their hosts. The compartmentalization of both symbionts into specialized bacteriocytes and their strictly vertical transmission may contribute to this convergence by imposing similar population bottlenecks and life-history dynamics. This pattern resembles the rate coupling recently described in psyllid nutritional symbioses (Degnan et al. 2025) but contrasts with the markedly different evolutionary rates reported for the dual symbionts *Nasuia* and *Sulcia* in leafhoppers (Vasquez and Bennett 2022), suggesting that tightly linked evolutionary trajectories may not always emerge from long-term co-obligate associations.

**Table 3:** Substitution rates reported in the literature for the primary aphid symbiont, *Buchnera aphidicola*. For each study, the dataset and methods used to estimate these average substitution rates are indicated.

| Study | substitution rate<br>(subst/site/Myr) | dataset | Calibration date | Substitution rate estimation<br>method | Interpretation |
| --- | --- | --- | --- | --- | --- |
| Brynnel *et al.* 1998 | dS ~0.0039–0.008 | Single protein-coding gene (*tuf*) from *B. aphidicola* from five aphid species | ~50-70 My (aphids from Aphidinae subfamily ) | Nb. of substitutions on the DNA fragment/ divergence dates | Long-term rate, sensitive to gene choice |
| Clark *et al.* 1999 | dN~ 0,0004-0,001<br>dS ~0,005-0,0095 | Multiple protein coding genes from *B. aphidicola* from four aphid species | ~50-70 My (aphids from Aphidinae and Pemphiginae subfamilies ) | Nb. of substitutions between pairs of species averaged across loci/divergence dates | Long-term rate, sensitive to gene choice |
| Tamas *et al.* 2002 | dN ~0,0016<br>dS~0,009 | Two *B. aphidicola* genomes (associated with *S. graminum*, *A. pisum*), estimation on 348 orthologous genes | 50 My (both aphids are from Aphidinae subfamily ) | Estimation of nb of substitutions/genome size/ divergence dates | Genome-wide long-term substitution rate, averaged across many loci |
| Gomez-Valero *et al.* 2007 | ~0.0043–0.0067 | Two introns and 1 pseudogene from *B. aphidicola* associated with seven aphid species | Divergence between Macrosiphini and Aphidini: 50 -70 My ago | Inference using estimation of the number of substitutions per site in each branch of a dated tree | Long-term neutral substitution rate; sensitive to gene selection |
| Moran *et al.* 2009 | ~0,11 | Nine *B. aphidicola* genomes associated with *A. pisum* | Date of introduction of the pea aphid in the USA (135 years) | Estimated nb. of substitutions / genome size/total tree length in years | Short-term substitution rate that reflects recent evolutionary dynamics within species |
| Perez-Brocal *et al.* 2011 | dN~0,0015 – 0,0036<br>dS~0,0036 – 0,008 | *B. aphidicola* genomes from four aphid species : 348 to 350 orthologous genes | Date of divergence of 112 My for aphid subfamilies and 50 My for the split between Macrosiphini and Aphidini | Estimated nb. of substitutions between genome pairs/ divergence dates | Genome-wide long-term substitution rate, averaged across many loci |
| This study | ~0,011 | *B. aphidicola* genomes from thirteen aphid species from the same genus (357 orthologous genes) | Divergence of a *Cinara* clade set at 25 My (secondary calibration estimated from a previous global phylogeny of *Cinara*) | Bayesian inference estimating evolutionary rates using Beast software (conducted on coding portions of the genomes) | Average long-term substitution rate, estimation using a phylogenetic method that models rate heterogeneity across the tree and date uncertainty |

Our study extends beyond evolutionary rates to investigate genome characteristics and signatures of selection in dual symbioses. In line with previous studies on *Buchnera* genome evolution (Chong, et al. 2019; Manzano-Marín, et al. 2023; Jousselin et al. 2024), we found a high degree of gene conservation within different *Buchnera*. Our investigation also shows that *Serratia* exhibits genome stability throughout its coevolution with aphids and *Buchnera*. Despite extensive genome reduction, its genome size and GC content remain largely unchanged across the clade, although gene losses continue to accumulate (Figure S4). Analyses of gene loss rates along the *Serratia* phylogeny actually reveal a heterogeneous tempo of genome erosion. While several bursts of gene loss are observed early in the phylogeny, consistent with rapid genome degradation following the transition from facultative to obligate symbiosis, more recent accelerations are also detected, notably along the lineage leading to *C. fornacula* and *C. mariana*. Together, these results support a scenario in which *Serratia* underwent substantial genome reduction early in its association with this aphid clade, followed by continued but uneven gene loss over evolutionary time. This pattern closely mirrors the evolutionary trajectories inferred for other obligate bacterial endosymbionts (Moran et al. 2008; Patino-Navarrete et al. 2013; Boscaro et al. 2017), suggesting that the mechanisms driving genome reduction are broadly shared across diverse endosymbiotic lineages.

It has been hypothesized that the presence of the co-obligate symbiont might accelerate the process of genome erosion in *Buchnera,* as it makes it redundant for several functions (Lamelas, et al. 2011). Similar hypotheses have been proposed for other co-obligate symbioses (Choi, et al. 2025a; Nozaki, et al. 2025; Michalik, et al. 2026). For instance the extensive pseudogeneization observed in *Walczuchella*, an obligate symbiont of giant scale insects, has been interpreted as a response to co-symbiont acquisition (Choi, et al. 2025a). The metabolic complementarity observed between *Buchnera* and *Serratia* in our study (Figure 1) is also consistent with a co-evolutionary process resembling the Black Queen Hypothesis, whereby redundant functions are progressively lost among interacting partners, promoting genome streamlining (Morris, et al. 2012). We found no evidence that redundancy between *Buchnera* and *Serratia* generally increases the evolutionary rate of shared host-provisioning genes, indicating that the persistence of redundant metabolic functions does not lead to a detectable overall relaxation of purifying selection. However, the correlation analysis revealed that while dN/dS values were significantly correlated between the two symbionts for genes from most functional categories, this relationship did not hold for host-provisioning genes. This lack of correlation suggests that homologous nutritional genes are no longer evolving under similar selective regimes in the two symbionts, potentially reflecting symbiont-specific relaxation of selection reflecting ongoing metabolic partitioning. This interpretation is consistent with the rapid functional specialization that follows the establishment of co-obligate symbiosis. Once overlapping nutritional pathways begin to partition between partners, the selective pressure acting on homologous genes may differ between symbionts. Consequently, redundancy may not increase the overall average evolutionary rate of shared nutritional genes, but instead lead to heterogeneous evolutionary trajectories. However, we have to remain cautious in our interpretation, as in a context of a drifting system the detection of selection signatures becomes very challenging. Furthermore as redundancy is rapidly eliminated following the establishment of co-obligacy (only a small number of host-provisioning genes are shared between *Buchnera* and *Serratia* Fig. 1), our analyses rely on very few genes. Systems retaining greater functional overlap, such as the co-obligate association between *Buchnera* and *Serratia symbiotica* in *Cinara tujafilina* (Manzano-Marín and Latorre 2014), may therefore provide a better opportunity to detect changes of selective constraints on redundant metabolic genes. Interestingly, genes involved in cellular processes and structures exhibited higher dN/dS values when shared by both symbionts than genes unique to either partner. This pattern may reflect host compensation for some cellular functions or the sharing of biological processes between symbionts.

Our intraspecific analyses of two *Cinara* species also provide a microevolutionary perspective on the parallel evolution of *Buchnera* and *Serratia*. We detected low levels of coding-sequence polymorphism in both *Buchnera* or *Serratia*, consistent with purifying selection, Strikingly, the amount of genetic variation were highly similar between symbionts, mirroring the comparable evolutionary rates inferred across the *Cinara* phylogeny. On the other hand, deletions accumulated in non-coding regions, particularly in *Serratia* genome, reflecting ongoing genome reduction. These mutational patterns actually closely resemble those previously reported for *Buchnera* within the pea aphid when occurring as the sole nutritional symbiont (Moran et al. 2009). Thus, despite its more recent transition to co-obligate status, *Serratia* can exhibit the same population-genetic signatures as the ancient primary symbiont *Buchnera.* Together, these results indicate that similar evolutionary forces act on both symbionts across timescales, from within-host polymorphism to long-term diversification. Comparable concordance between micro-and macroevolutionary patterns has been reported in leafhopper symbioses (Kwak et al. 2026).

## Conclusions

In conclusion, our study demonstrates that multipartite nutritional symbioses can remain evolutionarily stable over tens of millions of years, as evidenced by the parallel co-diversification of aphids and their two obligate symbionts. We describe a co-obligate di-symbiotic system in which *Buchnera* and *Serratia*, despite their distinct evolutionary origin, exhibit remarkably similar rates of molecular evolution and signatures of selection at both micro and macro-evolutionnary scales. These findings suggest that common selective pressures and demographic processes govern the evolution of both partners. The compartmentalization of symbionts into specialized bacteriocytes raises the possibility that host-mediated demographic conditions contribute to the convergence of their evolutionary trajectories. More broadly, our results support the view that once integrated into an obligate nutritional partnership, a newly acquired symbiont can rapidly adopt the evolutionary dynamics characteristic of ancient endosymbionts. Future investigations encompassing a more in-depth exploration of how the hosts can regulate symbiont densities as well as the roles of symbionts in host adaptation will further enhance our comprehension of these complex symbiotic systems. We expect such investigations, to answer why, in some lineages, the co-obligate symbiont, alongside *Buchnera,* has been repeatedly replaced (Manzano-Marín, et al. 2023) while we show here that it can be integrated as a long-term partner.

## Data and resource availability

The datasets and scripts underlying this article are available in Zenodo and can be accessed at 10.5281/zenodo.20026722 along with supplementary material. Available ENA accession numbers for mitogenomes, *Serratia* genomes and *Buchnera* genomes are indicated in Table S1 or to be announced, they have been submitted within the Umbrella BioProject PRJEB74807 Genome sequencing of the endosymbiotic consortia and mitochondria of Aphididae (CBGP - INRAE/UniVie).

## Conflict of interest

The authors have no competing interests.

## Supporting information

TableSupplementary

## Acknowledgements

We are grateful to the Genotoul bioinformatics platform Toulouse Midi-Pyrenees (Bioinfo Genotoul) for providing help and/or computing and/or storage resources. The authors are grateful to the CBGP-HPC computational platform. The funders had no role in study design, data collection and analysis, decision to publish, or preparation of the manuscript.

## Funding

This work was supported by the Marie-Curie AgreenSkills+ fellowship programme co-fundedby the EU’s Seventh Framework Programme (FP7-609398) to A.M.M. The Marie-Skłodowska-Curie H2020 Programme (H2020-MSCA-IF-2016) under grant agreement 746189 to E.J., the France Génomique National Infrastructure, funded as part ofthe *Inves-tissement d’Avenir* program managed by the *Agence Nationale pour la Recherche* (ANR-10-INBS-0009) to E.J., C.C., and V.B. JR was supported by an INRAe (Ecodiv) Phd Grant.

## Supplementary material

Annotated symbiont genomes, aphid mitochondrial genomes, data matrix used for phylogenetic analyses, ML trees as well as scripts used in this study are available in this zenodo archive in addition to the following supplementary material cited in the manuscript.

Table S1: Collection data and data accession numbers for *Cinara* samples analysed in this study. Each voucher number corresponds to a sample (i.e. an aphid colony).

Table S2: Assembly statistics and genomic features of sequenced *Buchnera* and *Serratia* strains associated with each aphid species. NCDS = number of CDS, Ncontigs = number of contigs, coding P =coding proportion.

Table S3: Tab S3: Genomic data from Buchnera aphidicola and Serratia sp used as outgroup. Mb= Megabases. Nb= Number.

Table S4: Table S4: History of gene loss events in Serratia associated with Clade A. List of CDS lost on each branch were inferred from the matrix of absence/presence of functional CDS.

Table S5: List of CDS in *Buchnera* genomes across species of clade A (1: CDS present, 0: CDS absent), dN/dS values estimated with Codml and results of selection tests using Busted, Codml (with various models), number of losses for each gene according to phylogeny.

Table S6: List of CDS in *Serratia* genomes across species of clade A (1: CDS present, 0: CDS absent), dN/dS values estimated with Codml and results of selection tests using with Busted, Codml (with various models), number of losses for each gene according to phylogeny.

Figure S1: Phylogeny of *Buchnera aphidicola* from *Cinara* spp. of Clade A based on a Bayesian analysis of a matrix of orthologous genes shared by all *Buchnera* strains, *Buchnera* from C*. tujafilina*, *C. cedri* and *Acyrtosiphon pisum* were used as outgroups. Asterisks at nodes stand for a posterior probability equal to 1. The genes lost in the different strains are indicated in red, dark red represents multiple losses and light red single losses. From left to right, the size of each *Buchnera* genomes is represented in light green in kb, dark green representing the coding proportion. The percentage of GC of each *Buchnera* is represented in light green.

Figure S2: Phylogeny of *Serratia symbiotica* from *Cinara* spp. of Clade A based on a Bayesian analysis of a matrix of orthologous genes shared by all *Serratia*. Co-obligate *Serratia* from C*. tujafilina*, *C. cedri* and facultative Serratia symbiotica from *Acyrtosiphon pisum* were used as outgroups. Asterisks at nodes stand for a posterior probability equal to 1. From left to right, *Serratia* genome sizes (in Mb) are represented in blue, dark blue representing the coding proportion. The percentage of GC of each *Serratia* is represented in light blue.

Figure S3: Correlation of root-to-tip branch lengths (in subs/site) from maximum-likelihood phylogenies of *Cinara* and *Serratia* reconstructed from matrices of shared genes only

Figure S4: Mapping of rates of gene losses per Myr in *Serratia symbiotica* associated with *Cinara* of Clade A on the calibrated phylogenetic tree of *Serratia*. Gene losses were computed from Table S4. To facilitate visualization and comparison across lineages, rates were log-transformed prior to graphical representation.

Figure S5: Histogram of the dNdS values depending of the status (shared or non shared between the two symbionts) and function of genes. Mean value are represented by dotted lines.

Figure S6: Scatter plot of log-transformed dN/dS values for orthologous genes shared between *Buchnera* and *Serratia*. Each point represents a single gene present in both genomes. Spearman correlation coefficient and p values are indicated for each gene functional category.

Figure S7: Intraspecific phylogenies based on maximum-likelihood analyses of *Serratia* (right) and *Buchnera* (left) genomes from *Cinara mariana*, showing congruent diversification patterns

Figure S8: Intraspecific phylogenies based on maximum-likelihood analyses of *Serratia* (right) and *Buchnera* (left) genomes from *Cinara fornacula*, showing congruent diversification patterns.

## Notes

https://zenodo.org/uploads/20026722

### Competing Interest Statement

The authors have declared no competing interest.

### Summary of Updates

After discussion with a colleague during a conference about this paper, we have slightly reworded the abstract and refined the correlation analysis of dn/ds. A supplementary figure has been updated also (figure S6). this is the last revsion before submission to a journal

